# Heterogeneity in Proline Hydroxylation of Fibrillar Collagens Observed by Mass Spectrometry

**DOI:** 10.1101/2021.04.12.439427

**Authors:** Michele Kirchner, Haiteng Deng, Yujia Xu

## Abstract

Collagen is the major protein in the extracellular matrix and plays vital roles in tissue development and function. Collagen is also one of the most processed proteins in its biosynthesis. The most prominent post-translational modification (PTM) of collagen is the hydroxylation of Pro residues in the Y-position of the characteristic (Gly-Xaa-Yaa) repeating amino acid sequence of a collagen triple helix. Recent studies using mass-spectrometry (MS) and tandem MS sequencing (MS/MS) have revealed unexpected hydroxylation of Pro residues in the X-positions (X-Hyp). The newly identified X-Hyp residues appear to be highly heterogeneous in location and percent occupancy. In order to understand the dynamic nature of the new X-Hyps and their potential impact on applications of MS and MS/MS for collagen research, we sampled four different collagen samples using standard MS and MS/MS techniques. We found considerable variations in the degree of PTMs of the same collagen from different organisms and/or tissues. The rat tail tendon type I collagen is particularly variable in terms of both over-hydroxylation of Pro in the X-position and under-hydroxylation of Pro in the Y-position. In contrast, only a few unexpected PTMs in collagens type I and type III from human placenta were observed. The reproducibility of the different sequencing efforts of the same sample is also limited especially when the modified species are present at a low population, presumably due to the unpredictable nature of the ionization process. Additionally, despite the heterogeneous preparation and sourcing, collagen samples from commercial sources do not show elevated variations in PTMs compared to samples prepared from a single tissue and/or organism. These findings will contribute to the growing body of information regarding the PTMs of collagen by MS technology, and culminate to a more comprehensive understanding of the extent and the functional roles of the PTMs of collagen.

**Abbreviations page:** Both the single letter and the three letter abbreviations of an amino acid will be used with the following additions: Hyp or O stands for 4R-hydroxylated proline and 3Hyp stands for 3- hydroxylated proline. When needed for clarity, the lower case single letter abbreviation will be used to represent the genomic DNA sequence, and upper case ones the sequence seen in the peptides.

## Introduction

The new technologies in mass spectrometry (MS) have transformed collagen research in recent years, and even expanded the field of collagen research to include archeology for the study of ancient species (1, 2). The high sensitivity of MS and tandem MS sequencing (MS/MS) also led to the identification of new post-translational modifications (PTMs) in fibrillar collagen (3–10). Fibrillar collagen is the major protein of bone, skin, cartilage, and blood vessel walls, and plays critical roles in many physiological and pathological events (11, 12). The newly discovered PTMs that are of particular interest is the 3-hydroxyproline residues (3Hyp, or 3O) in unexpected locations, since mutations in the enzymes involved in the formation of 3Hyp have been linked to severe cases of Osteogenesis Imperfecta (the brittle bone diseases) (13, 14). Yet, emerging from further studies of 3Hyp is an increasingly more heterogeneous pattern in terms of number, location, and the percent occupancy of this PTM (4, 10). Such varied patterns of 3Hyp make it challenging to pin down the specific molecular interactions involving 3Hyp. At the same time, collagens have been used as biomarkers for disease detection, species identification, and investigations of the involvement of collagen in cancer metastasis, tissue remodeling, the homeostasis of extracellular matrix and cell signaling, and the mineralization process of bones, to name just a few (15–18). The unpredictable nature of prolyl-3-hydroxylation has hampered the application of MS in these and other related research that rely on a precise knowledge of the genomic sequence and the PTMs of specific segments of collagen. Both the functional research of PTMs and the applications of MS for a broad range of quantitative studies of collagen depend on a comprehensive understanding of the extent and the variations of PTMs.

Collagen is a highly processed protein during its biosynthesis (11, 12). The gene products of collagen (often referred to as the α chains) usually comprise a triple helix domain together with propeptides at both the N- and C-termini. The triple helix domain of fibrillar collagens, which include collagens type I-III, V and XI, often contains more than 1000 amino acid residues in the uninterrupted (Gly-Xaa-Yaa) repeating amino acid sequence; while the Xaa and Yaa can be any amino acid. Collagen is known for its exceedingly high content of Pro: about 10-12% of the residues at each of the X- and Y-positions are Pro(19). In addition to the processing of the N- and C-propeptides, the residues of the triple helix domain are also heavily hydroxylated and glycosylated during the post-translational modification (20, 21). The major PTM of collagen is the prolyl-4-hydroxylation of Pro residues in the Y-position of the (Gly-Xaa-Yaa) triad (11).

This is a stable, invariant modification; nearly all Pro residues in the Y-position are hydroxylated to 4R-hydroxyproline (4Hyp). Some Lys residues in the Y-position are also hydrolyzed to hydroxylysine (Hyl), which are often glycosylated and/or form covalent cross-links through an oxidation process catalyzed by lysyl oxidase during tissue maturation (22, 23). Until recently, only one Pro residue in the X-position, Pro^986^ of the α1 chain of type I and type II collagen (the α1(I) chain and the α1(II) chain, respectively) was known to be a 3R-hydroxyproline (3Hyp), in which the hydroxyl group is appended on the *β*-carbon (the 3-position) of the pyrrole ring of the Pro instead of on the *γ*-carbon (the 4-position) of the pyrrole ring as in the case of 4Hyp (24). A more recent study using MS and MS/MS found several other Pro in the X-position of fibrillar collagens were also hydroxylated, including Pro^707^ of α2(I) chain, Pro^944^ of α1(II) and α2(V) chains, and Pro^470^ of α2(V) chain (9, 10). Some of the X-Hyps were later confirmed to be a 3Hyp by Edman sequencing. Multiple 3Hyps were also found in the repeating genomic sequence of gly-pro-pro (gpp) at the C-terminal end of the α1(I) and α2(I) chains, although their exact locations could not be specified (6). Differing from the 3Hyp^986^, which is invariant and has a nearly 100% occupancy, the newly discovered X-Hyp and 3Hyp residues are often found in a mixed population having the percentage of hydroxylated moiety ranging from 10% to 80% depending on the location, the type of collagen, the tissue, and the organism (4, 6, 9, 10).

The PTMs of collagen are considered to be essential for its secretion and self-assembly, and the immune responses of tissues (10, 20, 25–29), although much of the molecular mechanisms of their involvements remain unclear. Impaired prolyl-4-hydroxylation is the major cause of the condition of scurvy linked to the fragility in skin, blood vessels, and dentine (20). In this case, the tissue fragility was linked to the decreased stability of the collagen triple helix due to the lack of 4Hyp. Studies using triple helical peptides have firmly established the significant stabilizing effects of a 4Hyp in the Y-position compared to that of a Pro (30–33). Organisms having a higher physiological temperature often have a higher content of 4Hyp (34). The Hyl related glycosylation and cross-links were also considered an important part of the fibril stability, and the extent of the modification increases with the advance of the developmental stages(35–37). The understanding of the function of 3Hyp is more limited, except it is important for bone health (13, 38, 39).

Determining the biological functions of PTMs in fibrillar collagen is often confounded by the complex structural hierarchy of collagen fibrils (11, 12). The processed α chains of fibrillar collagen frequently have over 1000 amino acid residues. Three such α chains, they can be identical or different, come together in parallel to form the rod-shaped collagen triple helix, each about 300 nm in length. The Gly residues at every third position are packed at the center of the helix, while the side chains of the X and Y residues are largely exposed to solvent. The triple helix, however, is not the functional unit of collagen. The triple helices further self-associate laterally in a specific manner to form collagen fibrils having a unique 67 nm axially repeating structure known as the *D*-period. Any modifications of residues in X or Y positions can potentially impact the stability of the triple helix, the molecular recognition process during fibrillogenesis, and the interactions of collagen fibrils with cell receptors and other macromolecules during tissue development and function. Eyre and colleagues postulate the newly discovered 3Hyps are involved in fibrillogenesis because the locations of some of them are approximately a *D*-period apart (6, 8). However, considering the low occupancy at some of the locations, it remains to be evaluated at what extent of the hydroxylation the purported interactions involving 3Hyp will have a sustained impact on the fibril assembly. A systematic MS/MS characterization of rat type I collagen found an increased occupancy of 3Hyp with the developmental stages in rat-tail tendon, but the same study also reported a relatively constant extent of hydroxylation of type I collagen in bones and in skin (4). If the prevalent presence of 4Hyp is consistent with its structural role on the overall stability of the triple helix and the collagen fibrils, the highly diverged and sporadic presence of 3Hyp may suggest a more dynamic role for this unique collagen PTM, which may involve the hydroxylation of specific X-Pro residues at specific stages of development and/or in response to specific cues of the extracellular matrix. Similar dynamic PTMs were reported to be part of the ‘epigenetic code’ of histone and other proteins (40). Under such a dynamic regulation, 3Hyps are likely to present in low quantity: a few selected X-positions (comparing to more than 100 4Hyp residues in the Y-positions) in each α chain, and only in a subpopulation of the α chain. Both the locations of 3Hyp and the occupancy at a specific location may also vary depending on the physiological conditions of the tissues. Deciphering such dynamic patterns of 3-hydroxylation would require both robust protocols of quantitative detection and extensive sampling of collagens from different sources.

It is often difficult to delineate the variations of a PTM as part of the dynamic epigenetic regulation from the statistical nature of the techniques used for detection and/or for sample handling. MS and MS/MS are the technique of choice for studying PTMs of a protein, yet quantitative analysis of protein samples is often complicated (41, 42). The situation is particularly challenging for collagen due to the repetitive sequences and the high content of Pro residues. Fragmentation of Pro containing peptides is often inefficient due to the well characterized ‘proline effect’ in MS/MS which predicts a biased, sequence dependent potential to fragment N-terminal to a Pro bond during collision induced dissociation (CID) (3, 10, 43–45). In the case of the detection of 3Hyp the high frequency of the genomic sequence of pro-gly-pro-pro moiety further complicated the precise localization of the hydroxyl group without a good series of fragmented ions. Aside from functional dynamics, there is often an innate level of variations in the PTMs of a protein between individuals and between different organisms. Without a known priori on the statistical distribution of the PTMs in a specific tissue at a specific developmental stage, even a carefully designed study can only reflect a statistical snapshot. Collagens produced by recombinant systems may appear to be a well-controlled source of more homogeneous collagens. However, the expression of a foreign gene(s) can skew the PTM processes in a host cell leading to a different PTM pattern (2).

To gain a better understanding of the average impact of the prolyl-hydroxylation in both the X and the Y positions and the reliability of the detection by standard MS techniques, we carried out MS/MS sequencing of several samples of collagen from commercial sources and of collagen isolated from tissues. Our study revealed a more varied nature of the hydroxylation of proline residues in the type I collagen and substantial differences in the hydroxylation pattern among different collagens. The commercial collagens are often scorned as being impure because their productions often rely on batch collection of samples from mixed sources. However, this ‘mixed’ nature of commercial collagens can be a good statistical representation of the overall extent of different PTMs. Additionally, commercial collagens are very frequently used as standards for analytical analyses, and as extracellular matrix substitutes in various biological and biomedical studies that rely on specific interactions with residues on collagen including the PTMs. A better understanding of the PTMs of commercial collagen samples will enhance the sensitivity and reliability of these research efforts. Mapping out all the 3Hyp residues with the highest sensitivity and accuracy is not the main focus of this work. Rather, we seek to understand the reliability and reproducibility of the detection of unexpected hydroxylations using the standard MS approach. Furthermore, the uncertainties of PTMs on the X position complicate the fundamental premises of MS studies of collagen that assumes Pro in X-positions are unmodified, while those in the Y-position will inevitably have a mass increase of 16 due to the addition of the hydroxyl group. The finding of this work will, thus, contribute to both the understanding of the dynamics of 3Hyp and the applications of MS in other areas of collagen research.

## Materials and Methods

### Collagen preparation

Human collagen type III, human collagen type I, and rat collagen type I were purchased from Sigma. According to Sigma, human collagen type I and type III was purified from placenta, and rat collagen type I from tail tendon. The purchased collagens were solubilized in 20 mM acetic acid, pH 3, at 4 °C and 2.4 mg/mL. Fresh rat collagen type I was prepared from a single rat tail tendon following a procedure published by Dr. Sergey Leikin’s group, and the precipitated collagen was solubilized in 20 mM acetic acid, pH 3, at 4 °C and 3.0 mg/mL (46). Collagen was mixed with 5X SDS sample loading buffer containing 60 mM Tris-HCl pH 6.8, 25% glycerol, 2% SDS, 350 mM DTT, and 0.1% bromophenol blue, and was run on a 4-20% Precise gel or a 7.5% SDS PAGE gel and stained with coomassie blue. Bands of alpha chains were excised from the gels and submitted to The Rockefeller University for in-gel digestion and mass spectrometry analysis.

### In-gel Trypsin Digestion

The gel bands were reduced with dithiothreitol for 45 minutes at 55 °C, and alkylated with iodoacetamide for 30 minutes at room temperature in the dark. 10 μL of 0.02 μg/μL trypsin in 50 mM NH_4_HCO_3_/0.1% octyl glucopyranoside (OGP)/ 5 mM calcium chloride was used to digest each sample overnight at 37 °C in 50 mM NH_4_HCO_3_. The digestion was stopped with the addition of 5 μL of 10% acetic acid, and the peptides were extracted first with 30% ACN/5% TFA and then with 50% ACN/5% TFA. The samples were dried down in a Speed Vac to a few microliters.

### Trypsin In-Solution Digestion

The collagen solution was heat denatured and digested overnight with trypsin in 25 mM ammonium bicarbonate at 37 °C. The reaction was terminated by bringing the solution to a final concentration of 0.1% trifluoroacetic acid *MALDI-TOF MS Analysis* - Matrix, α-cyano-4-hydroxycinnamic acid, was prepared as a saturated solution in 50% acetonitrile/0.1% trifluoroacetic acid. Solution digests of collagen were spotted 1:1 with matrix onto a sample plate and allowed to dry. All spectra were acquired using a Voyager-DE STR mass spectrometer (PE Biosystems, Foster City, CA) equipped with a pulse nitrogen laser (λ =337 nm, 3 Hz frequency) in the reflectron positive ion, delayed extraction mode. Spectra from 100 individual laser shots were averaged.

### LC-MS/MS Analysis

The in-gel digests and solution digest samples were chromatographed using a C_18_ column on a Dionex HPLC eluded with a gradient of 0.1% formic acid and 100% ACN and introduced into a mass spectrometer. The sequencing was done at The Rockefeller University Proteomics Center.

### Data Base Search and Analysis

For peptide identification a Mascot (Matrix Science) search was performed. The .raw data files were converted to .dta or .mgf files which were then used to perform a search. The database User 0710 (in house database at Rockefeller Proteomic Center) containing human collagen type I and type III, and rat collagen type I, was used with oxidized methionine, proline and lysine as variable modifications. The Swiss Prot database was used for protein identification confirmation. For the freshly prepared rat tail, a summary report file was created using Discoverer version: 1.3.0.339, with the signal to noise threshold set to 1.5. Peaks6 software was also used. The reliability of the MS/MS largely depends on the detection of the fragmented ions. All the sequencing was determined using the Mascot search engine with a score of 40 or higher, and a probability of false identification of <10^-5^; an effort was made to manually check the identified ions in the sequencing outcome.

## Results

### 1. The variations in the hydroxylation of collagen

The heterogeneity of the hydroxylation of collagen α-chains can be observed at different levels by mass-spectrometry (MS). The sequencing by MS is carried out on trypsin digested peptides of collagen which usually range from 6-30 amino acid residues in length. The observed mass of one tryptic peptide that has undergone heterogeneous hydroxylation will resemble a mixture of species with mass differences of 16 (replacing –H by –OH) or multiples of 16. The presence of such mixed species was frequently observed in MALDI MS spectra of a total collagen digest (Fig. 1). The exact number of such peak-clusters varies among different collagens, and/or the different preparations of the same collagen. The observation of such peaks by MALDI MS depends on the trypsin digestion reaction and the signal level in the MALDI MS spectra. Only those peaks with the highest level of signal are labeled in Fig. 1; many others are present with low signal levels. Although it is not possible to identify these peptides by MALDI MS *per se*, given the widespread existence of peaks with mass variants of 16, it is unlikely for these peaks to be caused by the coincidental mass variations of different tryptic peptides. Rather, these peaks point to a mixed population of tryptic peptides with incomplete and/or ‘over’ hydroxylation of amino acid residues. The MS/MS sequencing study further supports this conclusion.

**Fig 1.**
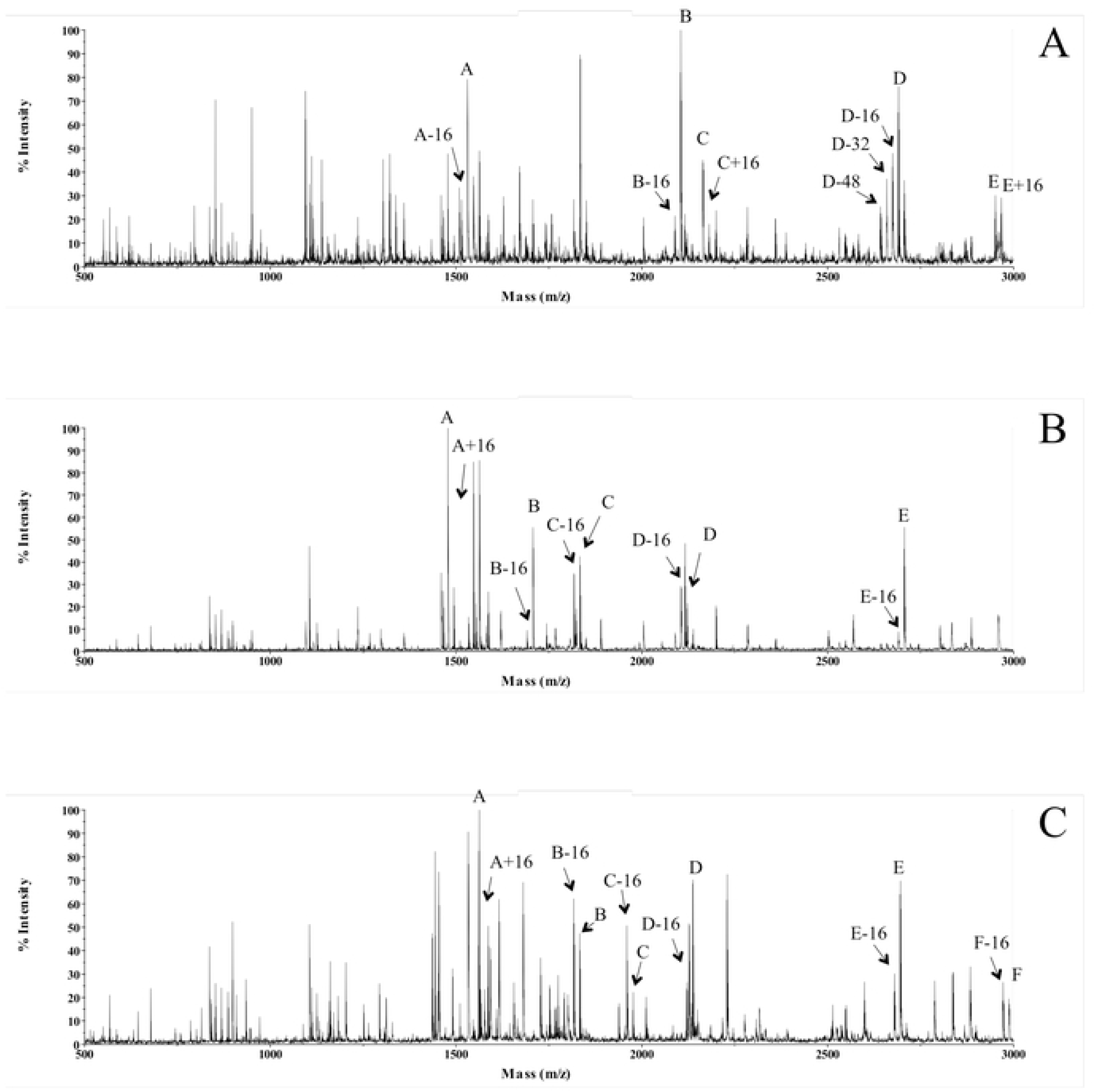
MALDI-TOF spectra of total trypsin digest. (A) Human collagen type III, (B) Human collagen type I, (C) Rat collagen type I. Peaks A – E in panel (A) and panel (B) and A-F in panel (C) are tryptic peptides of the corresponding collagens identified based on the agreement of their molecular weight (+1 ion) with that of the ‘theoretical value’ (assuming all Y-Pro as Hyp). The mass variants of 16 of each peak are labeled based on their mass differences from that of the ‘theoretical value’.

### 2. The O_x_ and the P_y_ residues identified by MS/MS sequencing of the collagens

For the clarity and the convenience of data presentation we will use O_x_ (or X-Hyp) and P_y_ (or Y-Pro), respectively, for a hydroxylated Pro in the X position and an unhydroxylated Pro in the Y-position to highlight the unusual hydroxylation results; the normal symbols of P (or Pro) and O (or Hyp) are used for Pro and 4-hydroxyproline in X and Y positions, respectively. Even by MS/MS sequencing, the exact modification often cannot be resolved with certainty with the mass information alone. In case a +16 mass is observed for a fragment of pro-gly-pro or pro-pro- gly sequence, for example, it is generally necessary to assume the hydroxylation of Pro is in the Y-position and not in the X-position in order to resolve the mass variations. Such fragments are common because of the high content of Pro in collagen and the proline-effect of MS/MS- sequencing (3, 10, 43–45). In compiling the sequencing data the ‘theoretical mass’ is calculated assuming *all* Pro residues in the Y-position are hydroxylated. Thus, a mass variation of -16 reflects an incomplete hydroxylation of Y-Pro residues, while that of +16 indicates an *additional* hydroxylation beyond the usual Hyp at Y-positions. In addition to Pro in an X position, a modified Lys (Hyl) in a Y-position or an oxidized Met (M_ox_) would also cause a mass change of +16 compared to their unmodified counterpart (47). A C-terminal Y-position Lys of the peptides can potentially be a hydroxylated Lys and contribute to the +16 mass variation. Considering the 7-fold decrease in trypsin susceptibility to Hyl relative to that of Lys, however, a C-terminal Hyl of a tryptic digest is an unlikely event (48). The C-terminal Lys residues are, thus, usually taken as one that is not hydroxylated with one exception: Hyl^87^ of the α2 chain of rat tail type I collagen (see later sections). The hydroxylation of this Lys was supported unambiguously by the +16 data of fragmented ions: all Y-Pro residues of this tryptic peptide are all fully occupied and there is no Pro in the X-position. The Lys in the equivalent position of human collagen was found to be hydroxylated by other methods (3). Hyl will not be seen in the sequencing results if glycosylated or cross-linked to neighboring peptides.

In the following, we only report the O_x_ and P_y_ residues that are unambiguously supported by a series of fragmented b- and/or y- ions. One typical example of unusual hydroxylation is given in Fig 2. There are 3 Pro residues in this 19-residue peptide from position 435-453 of the α_1_(I) chain of collagen isolated from a single rat tail tendon (srtt): two in the X-position (p^446^, p^449^) and one in Y-position (p^444^). The tryptic digest of this particular region of the α1(I) chain was partitioned into three populations with different hydroxylation. The sequencing outcome in Fig 2A reflects the expected hydroxylation pattern, with the p^444^ being hydroxylated and the mass of the peptide the expected value of 1680.76. The Hyp^444^ is supported by the identification of a nearly complete series of y- and b-ions and most directly by the y_10_ ion with a very strong signal. The sequencing outcome of the second population indicate the peptide carries a mass variant of +16 compared to the theoretical value (Fig. 2B), and the extra hydroxylation is unambiguously located on the p^449^ at the X-position, directly supported by the +16 values of y_5_ and y_6_ ions comparing to that in Fig 2A (*i.e*., the theoretical value). In the third case (Fig. 2C) the mass has a -16 variant, and again the strong y_10_ ion, as well as the -16 mass of the y_12_ and y_13_ ions demonstrated that the p^444^ at the Y-position was not hydroxylated. Thus, there are three different scenarios regarding Pro hydroxylation in this region: the one with expected Hyp^444^ in the Y-position, the one with O_x_, and the one with P_y_ . Since this particular collagen sample is isolated from a single rat tail tendon, the heterogeneous patterns of the hydroxylation reflects the natural variations of the Pro-hydroxylases and/or the variations accumulated over the development of this organism.

**Fig 2.**
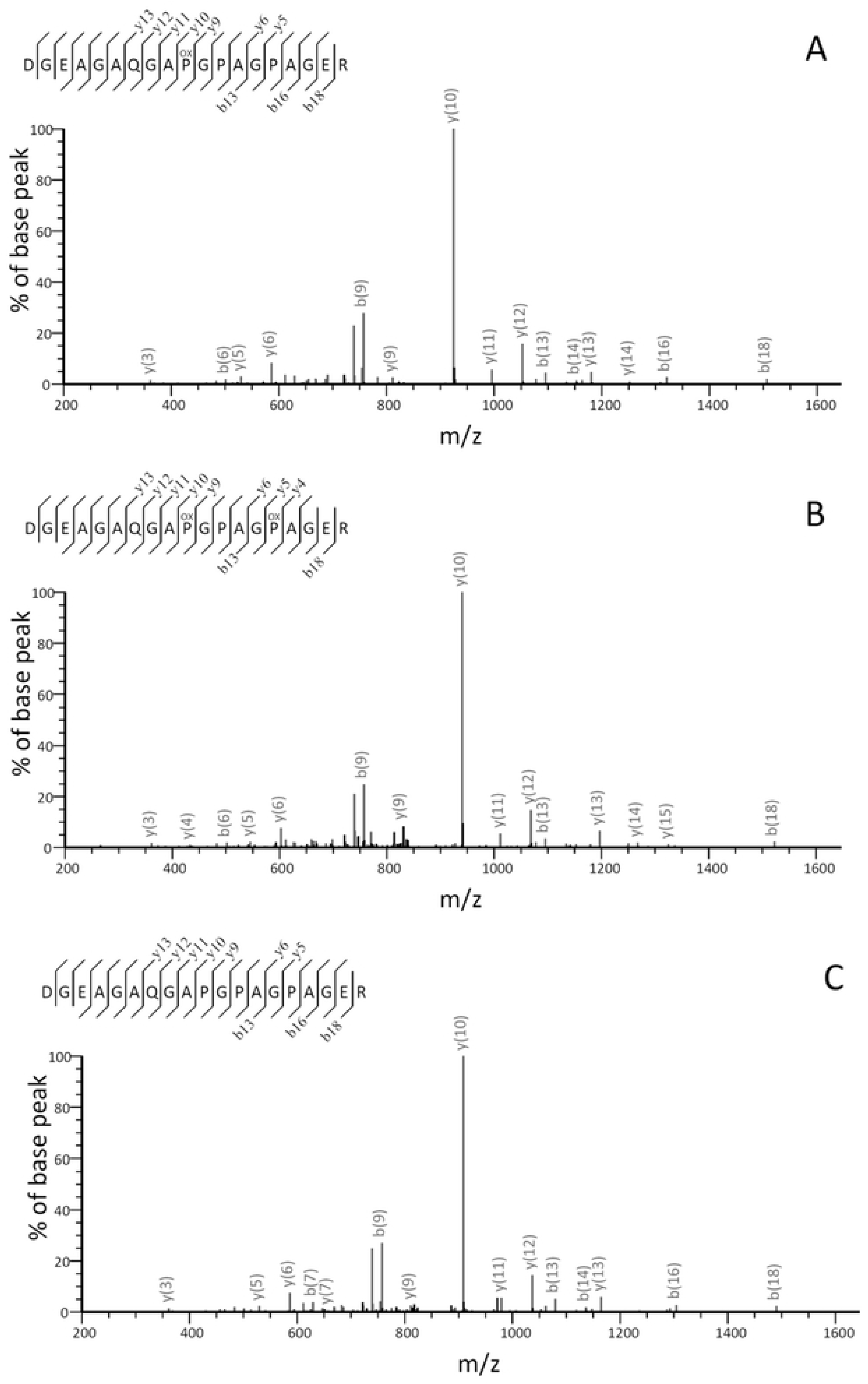
The MS/MS spectra of tryptic peptide ^435^dgeagaqgapgpagpager of rat tail tendon α1(I) chain. (lower case stands for the sequence from the genes). Sequencing outcome from ion 840.37762^+^ (1680.7551^+^) (upper panel), ion 848.37612^+^ (1696.7521^+^) (middle panel) and ion 832.38122^+^ (1664.7623^+^) (lower panel). The hydroxylation sites are shown as 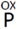. For clarity, only selected ions, those most relevant to the identification of residues are labeled.

The MS and MS/MS sequencing were carried out for the α1(I) and the α2(I) chains of type I collagen of rat tail tendon of a commercial sample (crt) and of a sample that is purified from a single rat tail tendon (srtt), the α1(I) and the α2(I) chains of a commercial sample of human type I collagen from placenta, and the α1(III) chain of human type III collagen from a commercial sample. The results of the peptides with unexpected hydroxylations for all five α chains of collagens are summarized in Tables 1-3 (details below); complete sequencing results of all collagen samples are given in the tables of the supplemental material S1. All the samples are sequenced at least twice in order to evaluate the reproducibility of the results. The reproducible findings are shown in boldface in the tables. In addition to the fragmentation data, the unusual hydroxylation of the peptides in Tables 1-3 is unequivocally identified by the variations of their masses (Δ*m*).

**Table 1.**
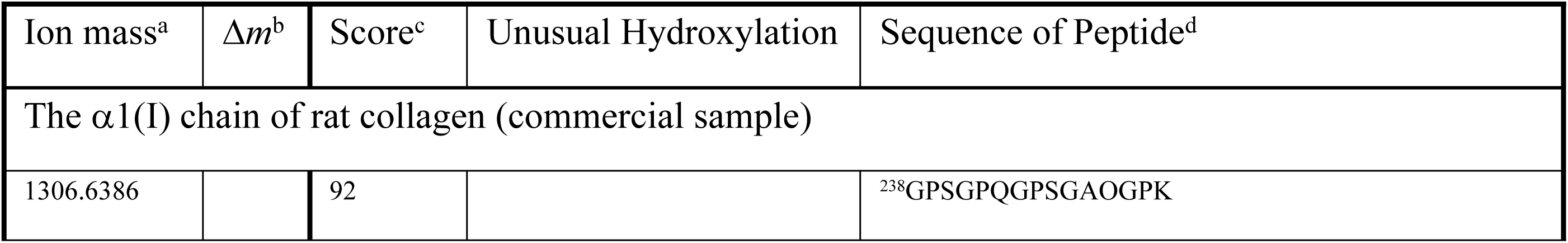
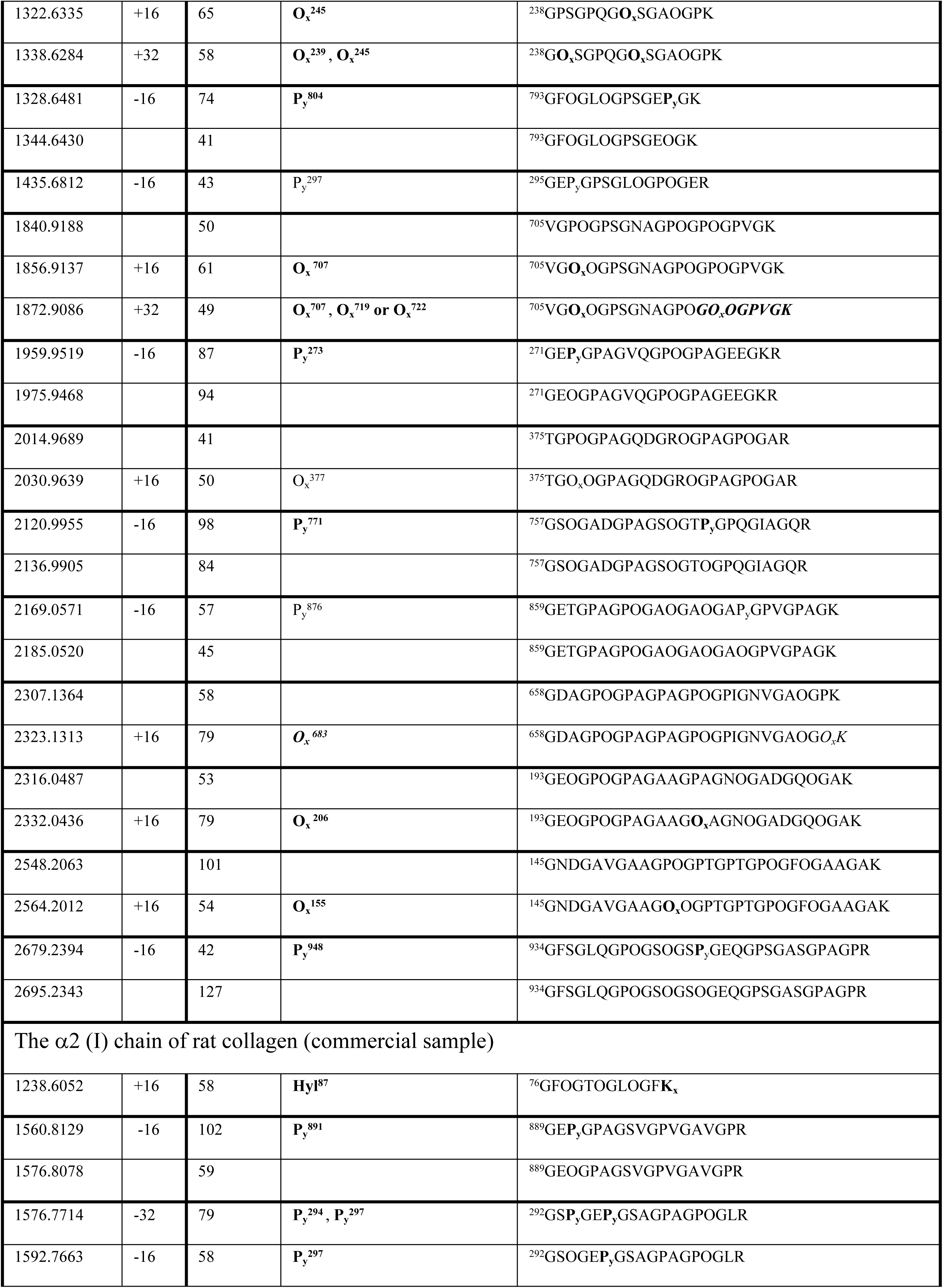
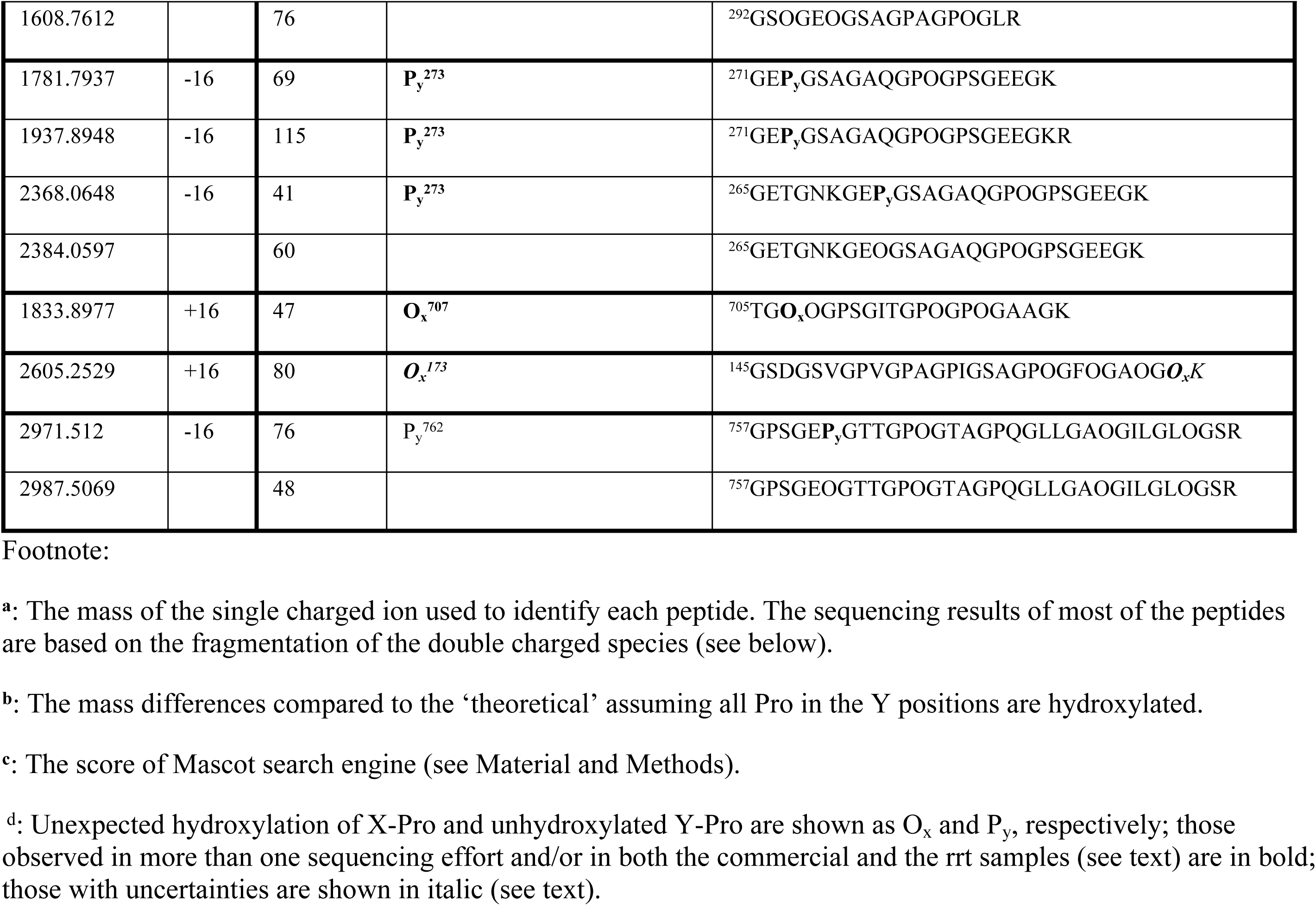
Peptides of rat type I collagen with mass variants of 16 (the commercial sample)

### 3. The over hydroxylation and under-hydroxylation of type I collagen from rat tail tendon

The results in Table 1 revealed a range of variations of the hydroxylation in both the α1 and the α2 chains of a commercial sample of the rat tail tendon type I collagen. The +16 or +32 mass confirmed the unexpected hydroxylations in 6 segments of the α1 chain at residues 145- 174, 193-219, 238-252, 375-396, 658-684, and 705-725; three over-hydroxylated segments were found in the α2 chain: at residues 76-87, 145-174 and 705-725. The hydroxylation sites of these peptides except that of residues 76-87 (peptide mass 1238.6052) are assigned to a specific X- Hyp based on the fragmentation ions; the +16 mass between residues 76-87 is assigned to the C-terminal Lys since it is the only residue in that peptide that can be hydroxylated. The precise location of the extra hydroxylation of peptide mass 2605.2529 (residues 145-174) of the α2(I) chain was difficult to resolve since the terminal pro-lys residues were not fragmented; both are the candidates for the hydroxylation (with a +16 mass). We are not aware of any other report on the hydroxylation of this particular Lys. Since Lys is more likely to be cleaved by trypsin than Hyl, we have tentatively assigned the hydroxylation site to be the Pro^173^ in the X-position, and the ***O_x_^173^***was shown in italic in Table 1 to highlight this uncertainty. Despite lacking a clear resolution of the location of the extra hydroxylation, the same peptide was observed more than once during multiple sequencing efforts as shown in the bold face in Table 1, which is presumably related to its measurable presence in this rat tail sample. The Lys residue in the equivalent position of the α1(I) chain in the peptide of residues 145-174 was resolved unambiguously, and for multiple times by the observation of the fragment having a terminal unhydroxylated Lys. This peptide, however, was found to have an over hydroxylation site on X-Pro^155^ (**O_x_^155^**). Similarly, for peptide mass 2323.1313 (residues 658-684) of α1(I) chain (Table 1), the ***O_x_^683^*** was assigned with uncertainty because the non-fragmented terminal Pro-Lys prevented an unambiguous assignment of the unexpected hydroxylation site.

In most cases a mixed population of variable hydroxylation was observed for a particular peptide. For example, the peptide with ion mass 1306.6386 (residues 238-252) coexists with the two over-hydroxylated species: a +16 species with mass 1322.6335 and a +32 species with mass 1338.6284 carrying, respectively, one and two extra hydroxylation sties. Due to the unpredictable and complex nature of the ionization process of MS and MS/MS, it is difficult to quantify the relative percentages of the various hydroxylated species by MS alone. In a few cases, such as for the α2 chain peptide with mass 2605.2529 (residues 145-174) only a single population with an extra hydroxylation was sequenced. Despite two sequencing efforts, a species with the theoretical mass was not observed. This lack of detection, however, does not rule out the existence of this population in the sample *per se*. This species may fail to be sequenced to an acceptable quality either due to poor ionization and/or fragmentation, or may have failed to be identified due to unexpected post-translational modifications. One limitation of mass-spec data interpretation is the inability to draw conclusions about peptides that are not selected for fragmentation.

Concurrently, incomplete hydroxylation was found in six and four regions, respectively, of the α1(I) and the α2(I) chains, located in the regions of residues 271-291, 295-309, 757-780, 793-806, 859-884 and 934-963 of the α1(I) chain, and of residues 271-290, 292-309 (having 2 different P_y_ residues), 757-789 and 889-906 of the α2(I). Remarkably, among the 11 detected P_y_ residues in both α chains, seven of them were found in the triplet GEP_y_.

One noticeable observation is the overlapping of the regions having unusual hydroxylation between the α1(I) and α2(I) chains. The region between residues 705 and 725 in both α chains contain multiple O_x_ residues: O_x_^707^ and O_x_^722^ in α1(I) chain and O_x_^707^ in α2(I) chain. Similarly, some of the P_y_ residues appear located in similar regions of both α chains as well: P_y_^273^ between residues 271-291, P_y_^294^ and P_y_^297^ between residues 295-309, P_y_^762^ and P_y_^771^ between residues 757-789. Combining the observations of different peptides from both α chains, the region of residues 271-309 in both α1(I) and α2(I) chains stands out as a particularly poorly hydroxylated region, missing two to three expected Hyp residues in the Y-positions of each α chain.

The finding of such a wide range of variations in hydroxylation of the type I chain is rather unexpected. The purity and the purification procedures of this commercial sample were called into question. In order to get a better understanding of the origin of the heterogeneity we purified the type I collagen from a single rat tail tendon (srtt). Interestingly, the sequencing result of this srtt sample turns out to be remarkably similar (Table 2). The 2 observed O_x_ residues in the α2(I) of the commercial sample and all but two (O_x_^206^ and O_x_^377^) in the α1(I) chain (Table 1) were reproduced in the srtt sample. Similarly, more than half of the P_y_ residues found in the commercial sample were also observed in the srtt sample. This srtt sample appeared to be particularly over-hydroxylated having 11 O_x_ residues in each α chain. The content of P_y_ is also higher: 10 and 9 P_y_ residues, respectively, were found in the α1(I) and α2(I) chains. The noticeably more heterogeneous hydroxylation pattern of this srtt sample, especially that for the α2(I) chain, may relate to the better overall sequencing outcome of the sample reflected, in part, by the better sequence coverage of this α2 chain (Fig. 3 and S1 Tables S1-S6). The identified O_x_ residues of the α1(I) chain of the srtt sample include the well-known 3-Hyp O_x_^986^; this section of the α1(I) chain of the commercial sample was not sequenced. Another interesting peptide with an additional Hyp of interest is the seven-residue peptide from the N-telopeptide region preceding the triple helical domain (ion mass 1452.7264). The Met residue in this fragment appears to be oxidized based on the detection of M_ox_ with neutral loss. In addition to the M_ox_, the Pro in the N-telopeptide appears to be hydroxylated (the P*). Since this Pro proceeds a Gly residue, which characterizes the canonical hydroxylation site of the prolyl-hydroxylase (C-P4H), its hydroxylation, although never reported before, probably does not come as too much of a surprise.

**Figure 3.**
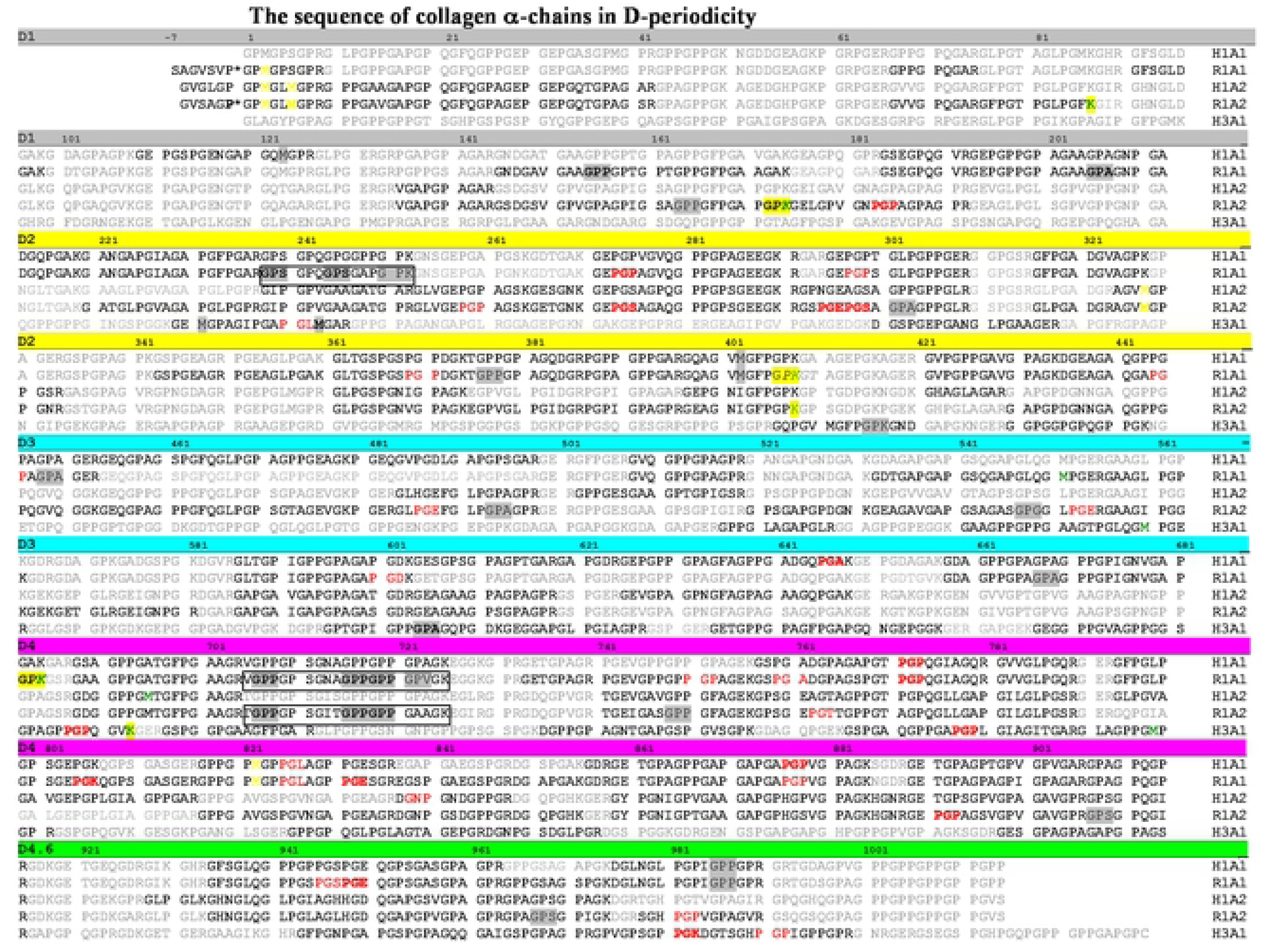
The mapping of individual unexpected hydroxylation sites on the α chains of collagen. Sequences of the α_1_(I) and α_2_(I) chains of human (H1A1 and H1A2, respectively), the α_1_(I) and the α_2_(I) chains of rat (R1A1 and R1A2, respectively), and the α1(III) chains of human (H3A1) were arranged by the *D*-periodicity according to Di Lullo *et al* (*49*): the 4 *D*- periods are highlighted by a colored bar of grey, yellow, cyan, and magenta, respectively; the 0.6 D is marked by the colored bar of green. The Gly-X-Y triplets including an O_x_ are in grey highlight. The P_y_ residues are lighted in red in the tripeptide unit of Y-Gly-X in order to reflect the potential connection with the enzyme selectivity of C-P4H (see text). The entire segment of the three highly variable regions (N- or C-HVR, see text) with multiple O_x_ residues are boxed. The hydroxylated proline in the telopeptide is P*. Hydroxylysines and the Gly-Pro-Lys tripeptide where the hydroxylation could not be precisely located between the Pro-Lys residues at the C- terminus of a peptide (see text) are highlighted in yellow with Lys in green font. The oxidized methionines are in yellow colored font. In all cases the PTMs observed from more than one detection/sample preparation were shown in bold font. Not sequenced regions are in faint grey. The amino acid sequence of the five collagen α chains were adapted from UniProt database.

The regions identified in the commercial sample where both α1(I) and α2(I) chains are over-hydroxylated, residues 238-252 and residues 705-725 (Table 1), also have multiple O_x_ in this srtt sample. The region of residues 705-725 revealed a particularly varied hydroxylation pattern having 1 to 3 O_x_ residues in both chains. By comparing to the commercial sample, another highly variable region stands out: residues 238-252 of α1(I) chain having up to 2 and 3 O_x_ residues in the commercial sample and the srtt sample, respectively. The over-hydroxylation, however, is not seen for the equivalent region of the α2(I) chain because of the non-homologous sequences: none of the equivalent X-residues of the α2(I) chain where an O_x_ is observed in the α1(I) chain is Pro. The poorly hydroxylated region of residues 271-309 is also under-hydroxylated in this srtt sample, lacking 1 and 3 expected Y-Hyps, respectively, in the α1(I) and the α2(I) chains. Other P_y_ residues that are frequently observed in both samples are P_y_^771^ and P_y_^948^ of the α1(I) chain, and P_y_^891^ of the α2(I) chain.

In fact, the sequenced peptides of the commercial sample in Table 1 appear almost as a subset of that included in Table 2A and 2B of the srtt sample. Thus, on this account, the commercial samples are quite representative of the averaged features of PTMs of collagens from the type I collagen of the rat tail tendon. All together combining the results of the two samples of type I collagen we have identified a total of 13 O_x_ in the α1 (I) chain and 11 in the α2(I) chains, also 13 and 10 P_y_ residues, respectively, in the α1(I) and α2(I) chains. The abnormal hydroxylation sites detected from *both* samples are mapped out on the sequences of the α-chains arranged in D-periodicity in Figure 3. The sites of O_x_ appear to be scattered rather uniformly throughout the α-chains except the two identified regions of highly variable hydroxylation patterns (HVRs): the N-terminal highly variable region (N-HVR) of residues 238-252 of the α1(I) chain, and the C-terminal highly variable region (C-HVR) of residues 705-725 of both α chains.

**Table 2A.**
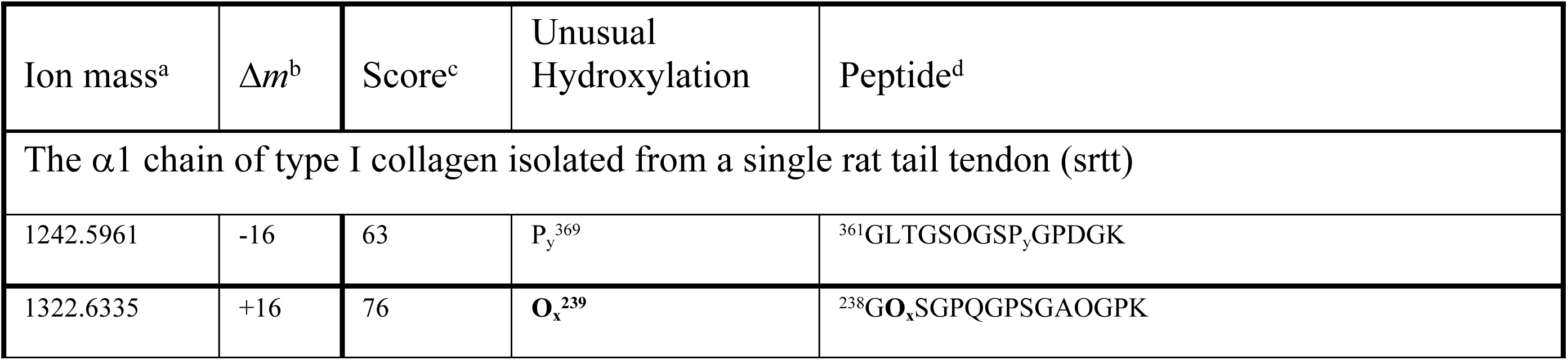
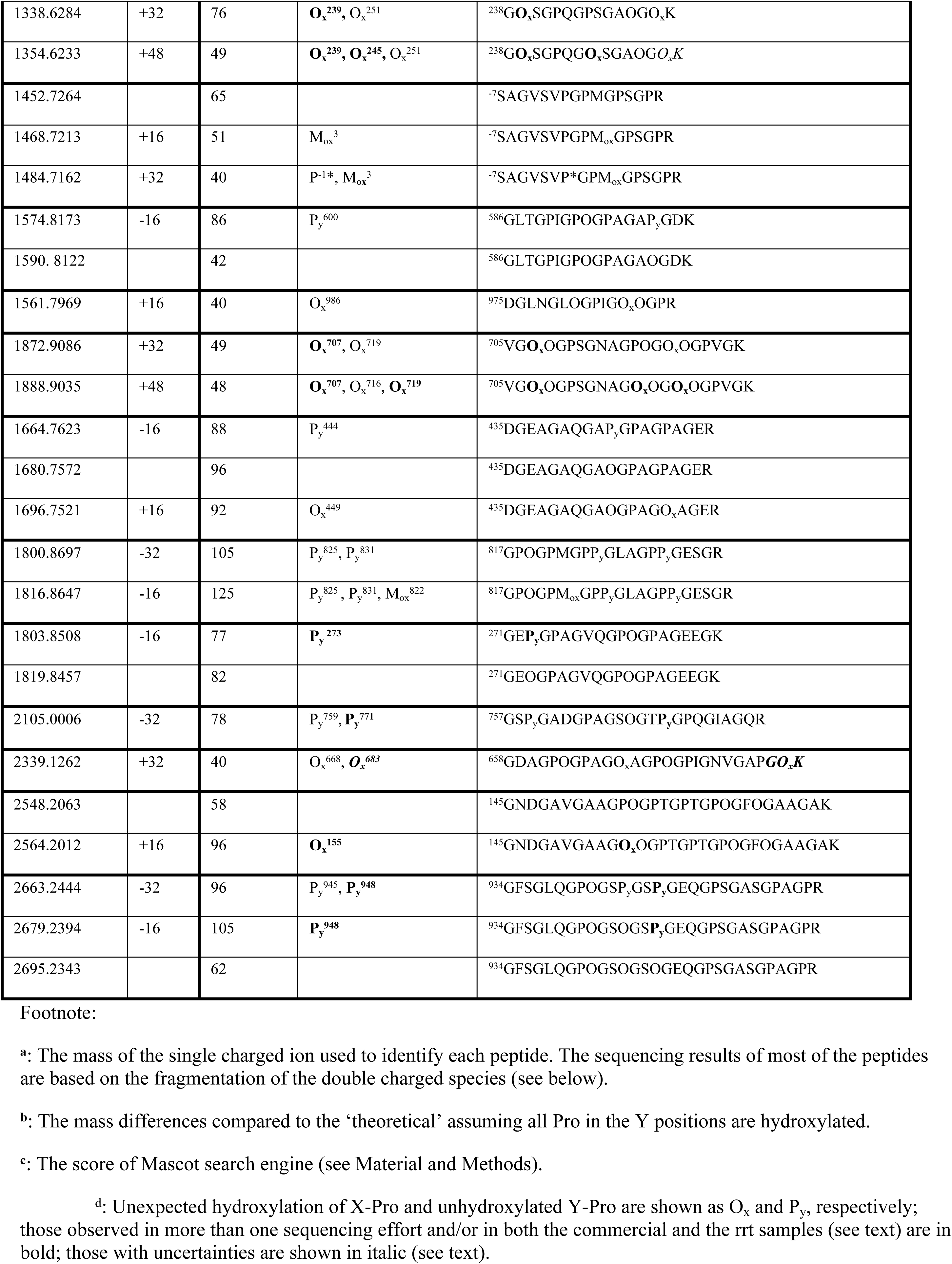
Peptides of type I collagen from a *single* rat tail tendon with mass variants of 16

**Table 2B.**
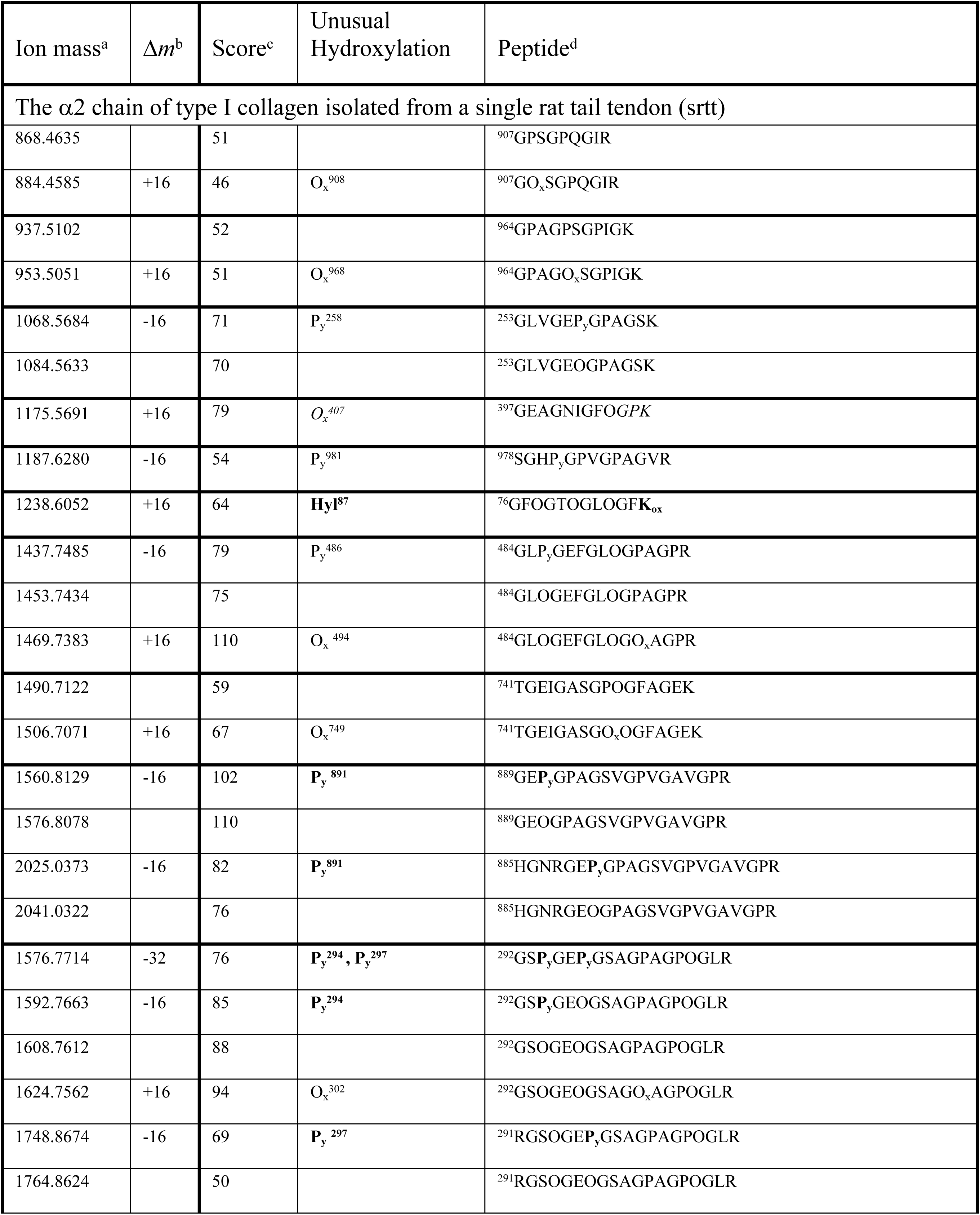
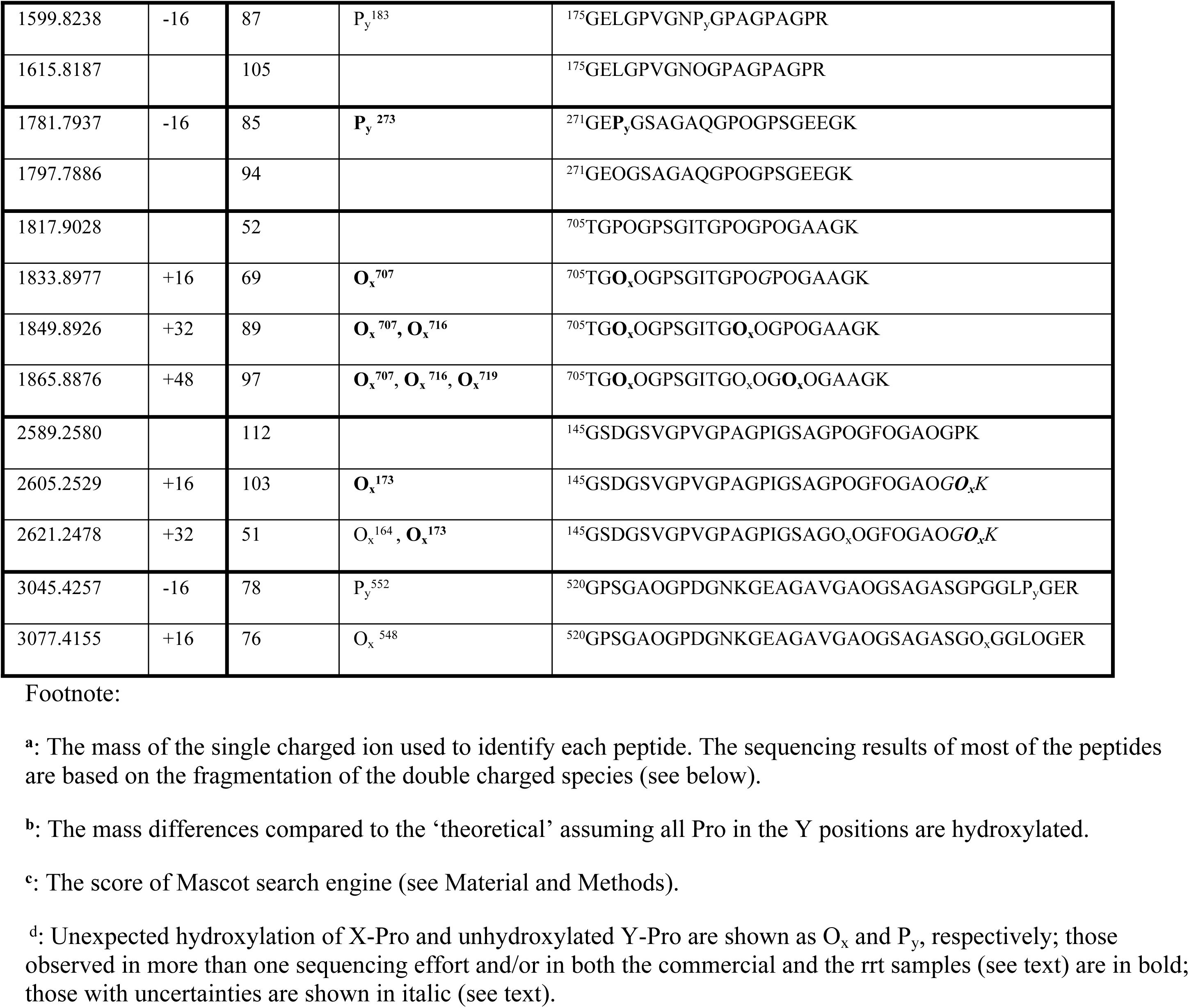
Peptides of type I collagen from a *single* rat tail tendon with mass variants of 16

The O_x_^155^ of the α1(I) chain is the only unexpected hydroxylation outside the HVRs that has been observed multiple times in both samples. The P_y_ residues also seem to cluster: in addition to residues 271-302 mentioned above, regions of residues 941-950 and 821-840 of the α1(I) chain, and 757-780 of both α1 and the α2 chains all have multiple P_y_ residues (Figure 3, Tables 1, 2A and 2B).

### 4. Human collagen type I and type III

The detected unusual hydroxylation sites of human collagen type I and type III are summarized in Table 3. Only the primary 3-Hyp at position 986 (O_x_^986^) of α1(I) was found as the X-Hyp in human type I collagen (Table 3A). Despite multiple sequencing efforts the peptide containing Pro^707^ of α2(I), one of the class-2 X-Hyp reported by Eyre and colleagues, was not sequenced. The Pro^707^ of α1(I) was sequenced but was found not hydroxylated in spite of the nearly identical amino acid sequences in this region between the two α-chains (Fig 3). The residue Met^822^ appears to be oxidized (Table 3A). The oxidation of Met is not a regular post-translational modification but an oxidation event usually found in cells under stress; it can also occur with sample handling (47, 50) (51). A few cases of incomplete hydroxylations were observed for both the α1(I) chain and the α2(I) chain. In general, the sequencing of the human type I collagen sample appeared to be rather clean, with only a low degree of unexpected modifications.

**Table 3.**
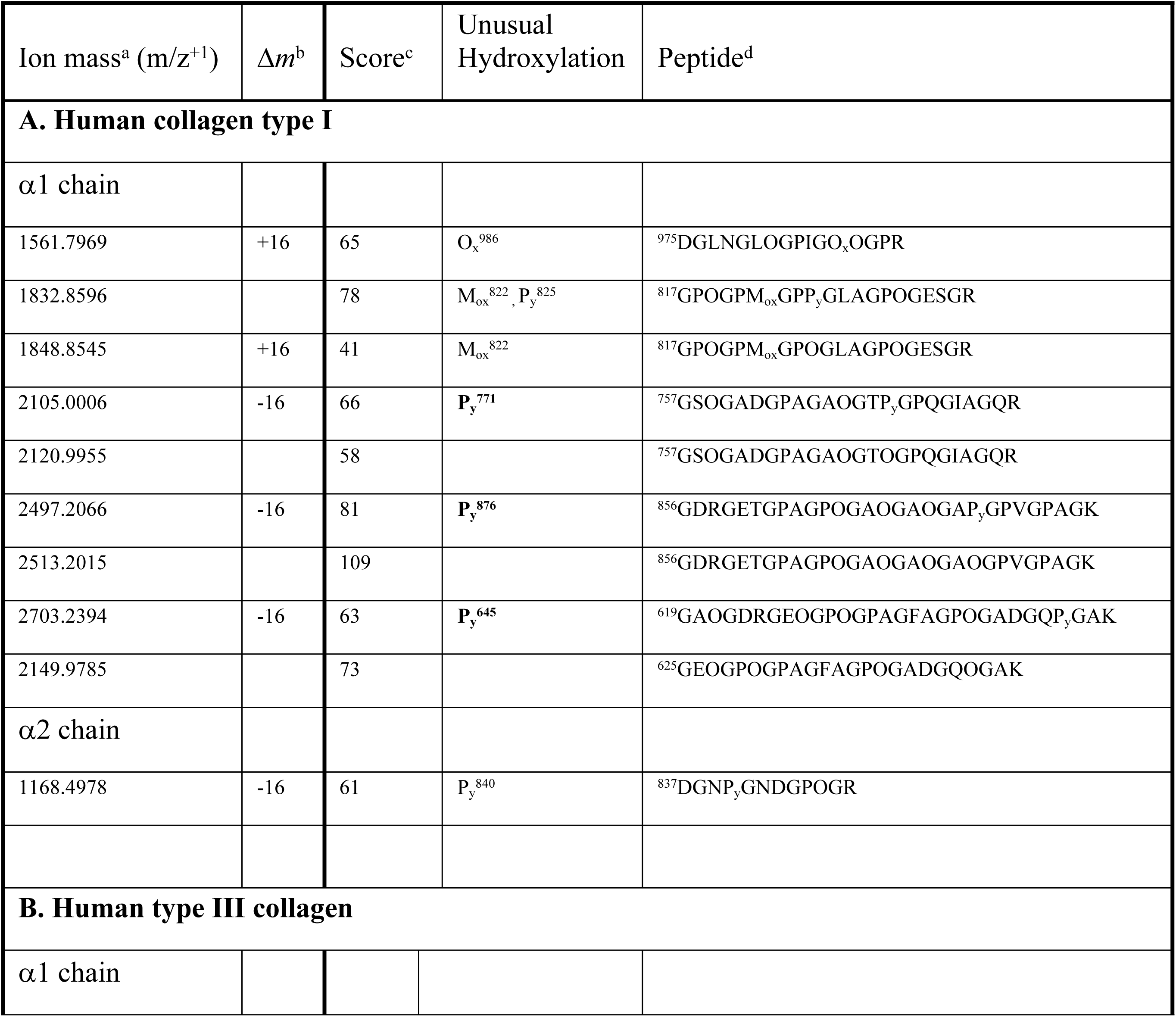
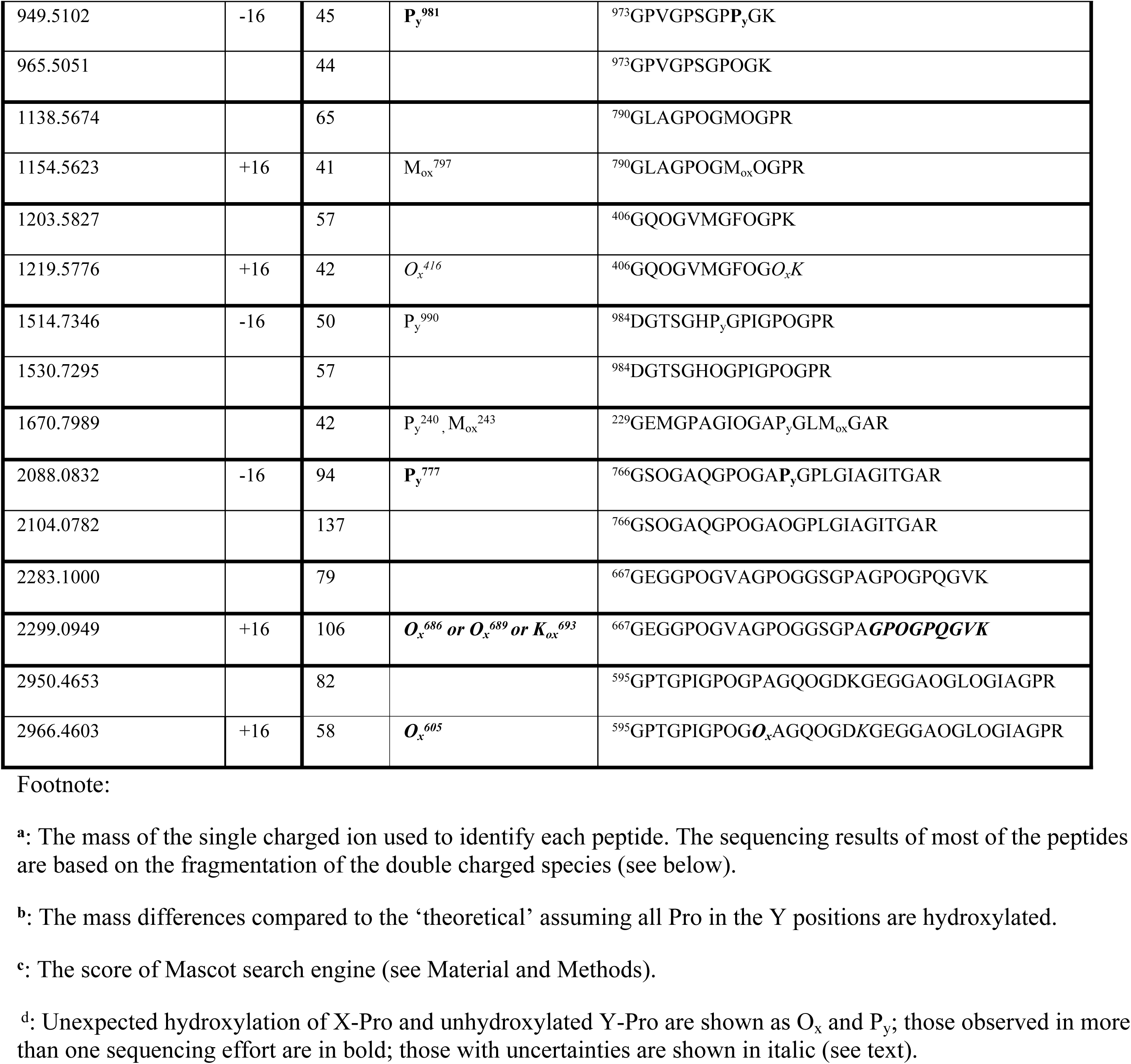
Peptides with mass variants of 16 in human collagen

Additional hydroxylation was seen in two regions of the α1(III) chain of the type III collagen (Table 3B): peptides of residues 406-417 (mass 1219.5776) and of residues 595-627 (peptide mass 2966.6403). The peptide with mass 2966.4603 contains an internal Lys^612^, the +16 mass was tentatively assigned to the hydroxylation of O_x_^605^ for lacking complete fragmentation between O_x_^605^ and Lys^612^. The peptide 406-417 (mass 1219.5776) carries the Met^412^, which can potentially be oxidized with a mass increase of 16; the fragmentation data has ruled out this possibility. Overall five P_y_ residues were detected in α1(III) samples, and the P_y_^981^ was seen with a very strong signal in every sequencing outcome.

In summary, the MS/MS sequencing results are mapped out on the sequences of the five α-chains arranged in *D*-periodicity in Fig. 3. Using the rather stringent sequencing criteria outlined in Materials and Methods the sequence coverage is about 35% for type III collagen and around 56% for the α_1_(I) and 46% for the α_2_(I) chains of human type I collagen, and about 62% and 67% for α_1_(I) and α_2_(I) chains, respectively, of rat tail tendon type I collagen combining both samples. Some observations of the O_x_ are consistent between the two different rat tail tendon samples, such as those in the HVRs; the others appeared sporadic. Most of the overhydroxylations seen in the rat tail tendon type I collagen are not present in human placental type I collagen. The consistent observations between the human type I collagen and that of the rat tail tendon include the Hyp^986^ of the α1 chain and P_y_ resides at positions 771 and 876 of the α1(I) chain.

## Discussion

By carrying out this study of the selected collagen samples we are hoping to gain a better understanding of the variations in the hydroxylation of fibrillar collagen in MS studies. Because of the high sensitivity of the MS and MS/MS approach, observing unusual hydroxylation of collagen proves to be a common event. Using a standard protein mass-spec sequencing technique we have detected unusual hydroxylation at several levels in rat type I collagen, human type I collagen and human type III collagen. The variations of the hydroxylation were supported by the spectrometry data for both the fragmented ions and the overall mass of the tryptic peptides. The over-hydroxylation was largely attributed to the hydroxylation of Pro in the X-positions, which is especially prevalent in both the α1(I) and the α2(I) chains of the type I collagen of rat tail tendon. The heterogeneity in the hydroxylation of human collagen type I and type III is much lower, reflecting the variations in enzyme selectivity of the hydroxylase among different species and/or tissues. As expected, most of the hydroxylated proline residues in the X-position are detected as a mixture; some may be present at a relatively low level, while others, as those in the highly variable regions (HRVs), are more prevalent and representative.

The repeated sequencing outcomes of the same collagen sample often carry high levels of variations as shown in Tables 1-3; detections of about half of the O_x_ and P_y_ residues are seen in multiple sequencing efforts (in boldface), while that of the others are less reproducible. In fact, the variations in the sequencing results of the two very different rat tail tendon samples are not in any way more substantial than that of the repeated sequencings of the same collagen samples.

Such varied outcomes reflected the complex and unpredictable nature of the ionization process of MS and sample handling (52). Each sequencing outcome often represents no more than a single sampling of a population consisting of heterogeneous modifications. The unpredictable ionization process is one of the major concerns for quantitative estimation of the populations of the sequenced peptides using MS/MS, especially when the sample is heterogeneous and the scope of the PTMs of the protein is not fully characterized. Sequence coverage will also affect the detection of PTMs, and this may be the reason that the canonical 3Hyp^986^ of the α1(I) chain was detected only once among multiple sequencing attempts of samples from human placenta and from srtt; this tryptic peptide containing position 986 was not sequenced at all in the commercial rat tail sample. Protocols using multiple proteases will result in a better sequence coverage, especially in cases like type III collagen where the tryptic peptides are often either too large or too small for reliable MS/MS results. On the other hand, despite the low sequence coverage, several unusual hydroxylations were observed rather consistently (Table 3, Fig 3). The observations of the O_x_ residues, in the highly variable regions of the rat tail tendon type I collagen is quite robust and consistent even among samples prepared from different sources.

The over hydroxylation observed in the C-HVR of the α1(I) and α2(I) chains of type I collagen in rat tail tendon is in keeping with the unusually high 3Hyp content of this collagen (53). The amino acid composition analysis estimated three to four 3Hyp in each of the α1(I) and the α2(I) chains of the rat tail tendon type I collagen, compared to only one 3Hyp, the Hyp^986^, in the rat α1(I) chain of type I collagen from bones or skin. The O_x_^707^, O_x_^716^ and O_x_^719^ of the α2(I) chains were subsequently identified as 3Hyp by N-terminal sequencing (4). The C-HVR also includes the location of the ‘class-2’ 3Hyp^707^ in the human α2(I) chain observed previously (9). Unfortunately this segment of the human α2(I) chain was not sequenced in our study despite repeated attempts; the same region in the human α1(I), which has the same amino acid sequence as that in the α2(I) chain, was sequenced, but no X-Hyp was found. The over-hydroxylation in the N-HVR of residue 238-252 of rat tendon α1(I) has never been reported before. Multiple O_x_ residues in this region in both samples of the rat tendon collagen were observed with reproducible results. Interestingly, the amino acid sequences of the two highly variable regions share limited homology; they are also rather different from the sequence surrounding the 3Hyp^986^. The two O_x_ residues of the N-HVR, O_x_^239^ and O_x_^245^ appeared in the peptide triad of GO_x_S, while the O_x_^707^ and O_x_^716^ of C-HVR in both α1(I) and α2(I) chains are in the more common GO_x_O moiety; but the O_x_^722^ of the α1(I) chain is in a GO_x_V triplet. The GOA and GOS are two moieties identified for X-Hyp of type V collagen (10). It is also worth noting that, similar to the 3Hyp residues identified by Eyre and colleagues, the two HVRs of rat tail tendon type I collagen are located exactly a 2*D*-period apart (Fig. 3), although the significance of it remains to be evaluated (9). No further effort was made to confirm the 3Hyp identity of the identified O_x_ in this study. While most of the newly identified X-Hyp residues have been confirmed to be 3Hyp, at least in one occasion an X-Hyp was later confirmed to be a 4R-Hyp (54).

The unhydroxylated Pro residues in the Y-position appear to be more common than X- Hyp among all α chains, with the highest content seen in the rat tail tendon type I collagen. Most of the detected Y-Pro residues are present as a mixed population having varied occupancies. Combining all the five α chains, the P_y_ residues were observed in 29 peptides. It is tempting to postulate the region of residues 273-302 of rat tail tendon type I collagen, where up to 5 P_y_ residues were found within a short stretch of 38-residues, to have unique conformational dynamics, since a Pro in the Y-position is known to significantly destabilize the triple helix compared to a Hyp (55). The real impact will, of course, depend on the percent of occupancy in these sites.

While incomplete hydroxylation has been known for some time, the site-specific data and the sequence motif of the missed-hydroxylations have not been reported before. The sequence information of the P_y_ residues may relate to the substrate selectivity of the prolyl-4-hydroxylase (C-P4H). Studies using short peptides established that the enzyme recognizes Pro-Gly-Xaa triplets during hydroxylation, where the Pro is the residue to be hydroxylated, and the selection of the Pro in a Y-position is affected by the conformation around the -Gly-Xaa residues (56–58). The hydroxylation takes place on the nascent polypeptide chains before the formation of the triple helix. Despite the higher than normal content of the Pro residues, the unfolded α-chains of collagen are not known to assume any well-defined conformation, although isolated segments may temporarily adapt to polyproline II (PP II) like or *β*-turn like *ϕ* and *φ* angles. Specifically, the type II *β*-turn bent between the -Gly-Xaa was considered to favor the binding of C-P4H and thus, the hydroxylation of the Pro proceeding the Gly; while the PPII conformation in -Gly-Xaa was considered inhibitory (57, 58). Residues Ala, Leu, Ile, and Phe in the position of Xaa were found to favor a β-turn around the Gly, and a Pro favors a PPII *ϕ* and *φ* angles (58). Our finding here appears to reflect this conformational preference of C-P4H in vivo. If we consider the P_y_ as a *miss* of the C-P4H in selecting a Pro in a Y-position for hydroxylation, the hydroxylation action appears to be particularly slippery in the sequence context of Pro-Gly-Pro. Nineteen out of the 36 P_y_ residues identified in the 5 α chains are in the P_y_-Gly-Pro moiety. On the other hand, this high occurrence of P_y_-Gly-Pro moiety may simply reflect the higher frequencies of genomic sequence pro-gly-pro in fibrillar collagen. The other frequent *misses* include P_y_ in Pro-Gly-Glu (5/33), Pro-Gly-Ser (4/30) triplets, and Pro-Gly-Leu (3/33) triplet. Among the identified P_y_ residues, the P_y_^771^ of α1(I) in a P_y_-Gly-Pro moiety is the only one that is identified in both human and the two rat tail tendon samples. The P_y_^876^, also in P_y_-Gly-Pro moiety, of α1(I) was detected in human and the commercial sample of rat tail tendon, but not in the srtt sample. There also appears to be an overrepresentation of P_y_ residues in a GEP_y_ tripeptide in rat type I collagen: 8 out of the 23 identified P_y_ residues are in a GEP moiety. It is not clear if the Glu preceding the Pro in the Y-position affects the selectivity of C-P4H in rat tail tendon. Other more common sequence motifs for P_y_ are GAP_y_, GPP_y_ and GSP_y_. It is also unclear if the missed hydroxylation of these residues has any functional roles for collagen.

## Conclusions

The high sensitivity of MS/MS sequencing has revealed a subpopulation of collagen that bares unexpected hydroxylated Pro in the X-positions, and unhydroxylated Pro in the Y-positions. The detection of X-Hyp and the Y-Pro by mass-spec sequencing is impacted by both the tissue/organism dependent variations of the Pro-hydroxylation reaction and the statistic nature of the technique. The detection of some modifications, however, is rather robust and can be used as a biomarker for general applications using MS/MS. A more thorough understanding of the dynamics of the specific PTM of 3-Hyp in the X-position and its role in epigenetic regulation will require a knowledge base that is broad enough to reflect the statistical nature of both the variations of the PTMs in different tissues and organisms, and the reproducibility of their detections by MS/MS.

## Acknowledgements

The authors are indebted to Dr. Sergey Leikin, and Dr. Elena N. Makareeva for their help preparing rat tail collagen, Dr. Milica Tesic Marks for some sequencing and data processing, Dr. Henrik Molina for Discoverer software assistance and the overall guidance on mass-spec technology. We would also like to thank Drs. Rebecca Strawn, James San Antonio and late Adel Bosky for critical reading of this and a previous version of the manuscript.

## Supporting Information

S1: The Complete Sequencing Results

